# Histone modifications at the transcription start site of non-coding RNA revealed regulatory patterns of activation in *Caenorhabditis elegans* embryo

**DOI:** 10.1101/2020.09.24.311308

**Authors:** Megan A. Bandeira, Max E. Boeck

## Abstract

Histone modifications play an essential role in regulating recruitment of RNA polymerase II and through this regulation of transcription itself. Which modifications are essential for regulating the transcription of non-coding RNA (ncRNA) species and how these patterns differ between the different types of ncRNA remains less studied compared to mRNA. We performed a principal component analysis (PCA) of histone modifications patterns surrounding the transcription start site (TSS) of ncRNA in an attempt to understand how histone modifications predict polymerase recruitment and transcription of ncRNA in early *C. elegans* development We found that our first PCA axis was a better predictor of polymerase recruitment and expression than any single histone modification for ncRNA and miRNA. This indicates an integrated analysis of many histone modifications is essential for predicting expression based on histone modifications and that each ncRNA species have unique regulation of RNA polymerase recruitment through histone modifications.

## Introduction

Nucleosomes are modified across the genome in response to a variety of transcriptional needs. These histones are actively modified with residues that affect how tightly the DNA is wrapped (1). For example, the addition of an acetyl group to the histone is an indicator of active transcription whereas histone methylation, or the addition of a methyl group can be an indicator of repressed transcription, especially at the transcription start site (TSS). Understanding the exact pattern of modifications needed for proper transcriptional activation and repression is essential to understanding everything from basic biology to disease phenotypes like cancer and diabetes (2).

Non-coding RNAs regulate gene expression and chromatin structure by binding to RNA and disrupting their stability and by binding proteins (3). Different species of ncRNAs regulate different events in the cell. We focussed on ncRNAs designated as miRNA, tRNAs, snRNAs, snoRNAs and general ncRNAs in the *C. elegans* genome. microRNAs (miRNA) which gene expression by targeting 3’UTR of mRNA leading to degradation or inhibition of translation (4). Transfer RNAs (tRNA) play important roles in protein synthesis. Small nuclear RNA (snRNA) plays a role in splicing mRNA as well as polyadenylation and mRNA stability (5). Small nucleolar RNAs (snoRNA) are responsible for carrying out site-specific base modifications of other ncRNAS (6). ncRNAs regulate gene expression through post transcriptional regulation. ncRNAs interact with other RNAs, DNA, and proteins and are involved in regulating which genes are translated into proteins as well as having important roles in several cellular processes including intracellular communication and protein synthesis (7). Predicting target expression of these ncRNA species, then, would be vital to our ability to understand essential cellular processes (8).

Different histone modifications are known to signal different transcriptional states. Acetylation of histones is almost always associated with active transcription with H3K27ac known to precede actively transcribed protein coding genes (9) while H4K8ac is thought to mark transcriptional elongation (10). Meanwhile histone methylation is more complicated; while marks such as H3K4me3 and H3K79me2 are associated with actively transcribed genes H3K27me3 is associated with repressed transcription (10, 11). Histone modifications also play an essential role in recruiting RNA polymerase 2 to transcription start sites. (12).

## Methods

### Datasets

Data on expression levels for non-coding RNAs during development in *C. elegans* as well as ChIP-seq data for histone modification signals were extracted from existing datasets (13, 14). We based both our expression and our annotation of ncRNA species and location on the Boeck et al. dataset. The average signal for each histone modification was calculated at different points around the transcription start site (TSS) for all annotated ncRNA species using cistrome sitepro for the following histone modifications: H3K20me1, H3K27ac, H3K27me3, H3K36me3, H3K39me1, H3K4me1, H3K4me2, H3K4me3, H3K79me2, H3K9me3, H4K8ac, HTZ1 and nucleosome positioning using MNase-seq data (designated dyads in our analysis) (13, 15). Total signal was extracted 1000 bp upstream and downstream of the TSS using 10bp bins. The 50bp surrounding the TSS was used to create our principal component analysis (PCA) of histone modification (16). Signal 1000bp surrounding the TSS was used to create composite histone signal surrounding ncRNA TSS.

### PCA Creation

The histone modification data were separated by ncRNA species and analyzed separately. The correlation of the vector of the PCA axes with ncRNA expression was then calculated. The first and second principal components for each PCA were plotted into a biplot. Additionally, ncRNA species were grouped into quantiles based on expression levels with higher expression levels in the first quartile and those with lowest expression levels in the fourth quartile. Ellipses of each quartile were plotted. Similar plots were made for RNA polymerase quartiles.

### Binning on PCA Axis 1

Each ncRNA species were then binned into quartiles based on their score in the first principal component with ncRNA species with the highest PCA score in the first quartile and those with the lowest PCA score in the fourth quartile. These were then plotted by expression level into a box plot. Average histone modification signal was plotted for each quartile for each ncRNA species. Composite histone modification signals at and surrounding the TSS of ncRNA species in each of these bins were then plotted for each of the four most enriched histones based on signal.

## Results

### PCA Model Generation

Based on the Principal Component Analysis and subsequent biplots (Fig 1), we find that the histone modifications H3K36me3 and H3K79me2 contribute strongly to the first principal component while the modifications H3K4me2, H3K20me1, and dyads contribute strongly to the second principal component for all small RNAs. We binned the different ncRNAs species based on expression and plotted them on a biplot of the PCAs we generated. When we plotted the PCA biplot with RNA expression bins, we found evidence of expression bins clustering along the PCA1 axes for miRNA, ncRNA and snRNA, but not for tRNA and snoRNA. This indicates the PCA is clustering the RNAs based on expression levels.

**Figure 1.**
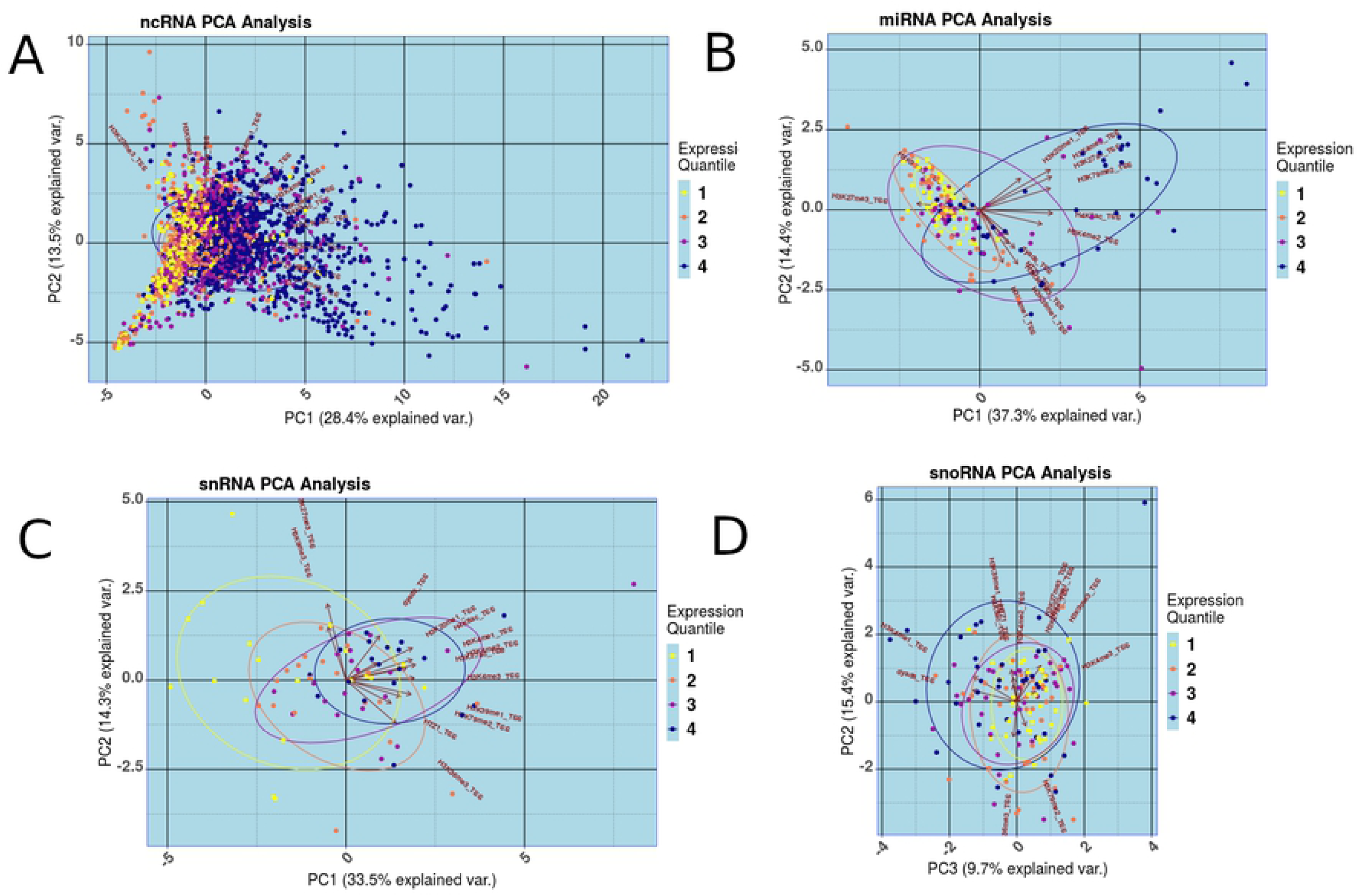
PCA plots of Histone Modification Signal Surrounding the Transcription Start Site of ncRNA, miRNA, snoRNA and snRNA. TSS are colored by which expression quantile the RNA species falls in. The first quantile contains RNA with the highest expression levels while the fourth qualtine contains RNA with the lowest expression levels. Quantiles are colored yellow, orange, purple, blue with yellow the highest expression quantile and blue the lowest A - ncRNA PCA plot, B - miRNA PCA plot, C - snoRNA PCA plot, D - snRNA PCA plot.

### PCA prediction of Transcription and RNA Polymerase Recruitment

We next calculated the correlation of our PCA with the expression for the different ncRNA species as well as the RNA polymerase 2 (RNAP2) signal at the TSS and compared this correlation to the correlation of individual histone modifications with each ncRNA species’s expression and RNAP2 signal (Fig 2A). We found strong correlation between histone modification signals and expression of miRNA, snRNA, and ncRNA.The first principal component showed relatively high correlation with most ncRNA species and had a higher correlation with expression than any single histone modification for both ncRNA and tRNA and all but one histone modification for miRNA and snRNA. This indicates that the PCA can predict expression to a relatively high degree. We next compared how well our PCA predicted RNAP2 surrounding the TSS of each of the ncRNA species (Fig 2B). We found that our PCA had higher correlation with RNAP2 surrounding the TSS for miRNA, ncRNA and tRNA than any single histone modification. Because pol2 is a strong indicator of transcription, this indicates that the principal component is a strong model to predict transcriptional activation of these RNA.

**Figure 2.**
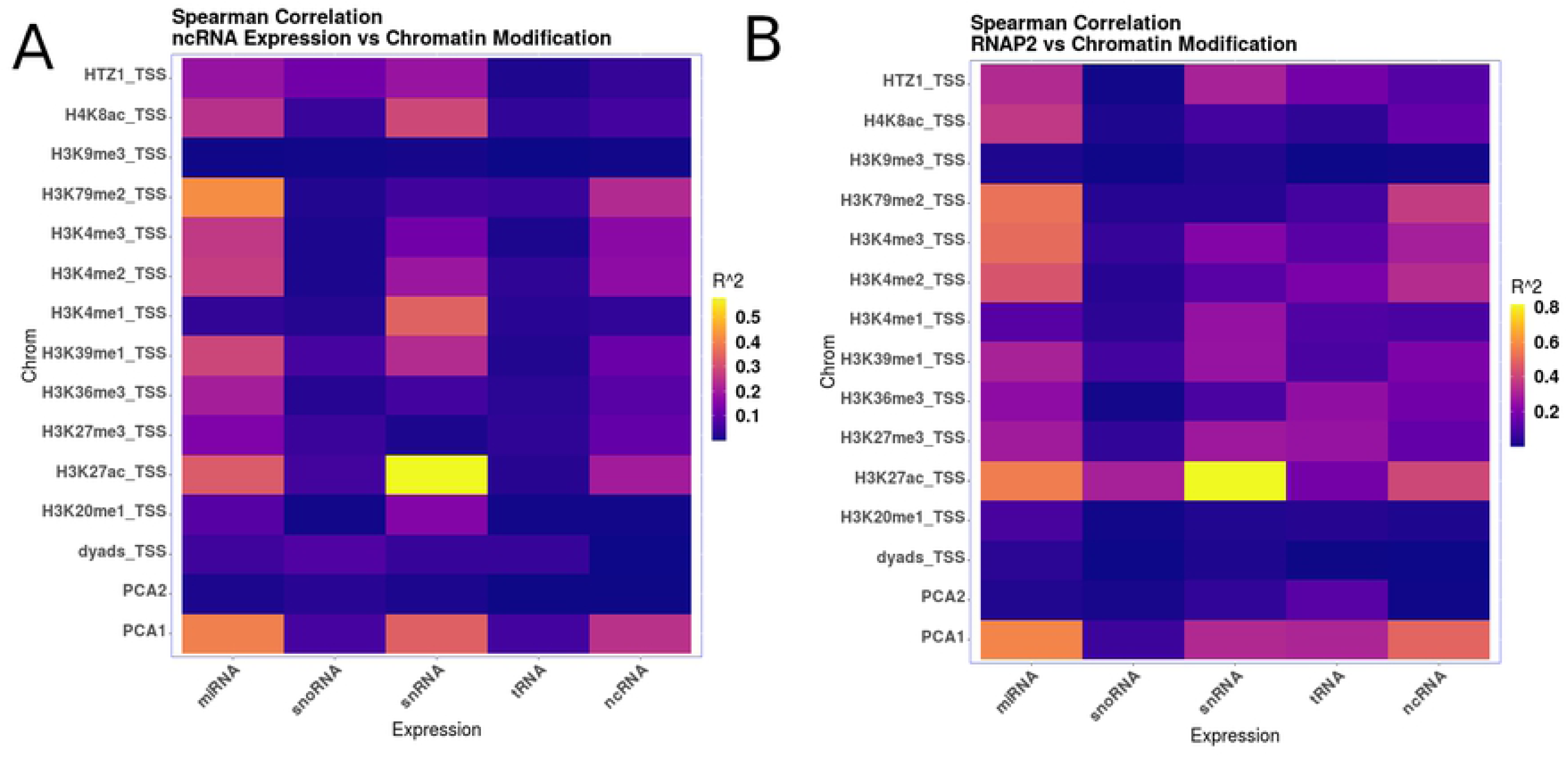
Spearman Correlation of Expression and RNA Polymerase 2 binding with all the histone modifications used to generate our PCA model along with PCA1 and PCA2 of the model. R squared is plotted for each comparison with yellow showing high correlation, purple medium correlation and blue no correlation. A - Correlation of expression with histone modification and PCA axes. B - RNA polymerase binding at the TSS correlated with histone modification and PCA axes.

We binned our ncRNA species into four bins based on the PCA1 axis and plotted both expression and RNAP2 signal (Fig 3 and 4). We found that there was a strong trend for higher expression and higher RNA pol2 signal dependent on PCA1 for miRNA, ncRNA and snRNA. Similar to the correlation calculations, this trend was more pronounced for RNAP2 binding. Expression levels of ncRNAs increased significantly as the PCA1 score decreased (Fig. 3A). The expression of miRNA increased as PCA1 score decreased (Fig. 3B). This is also true for snRNA expression (Fig 3D). This indicates that the PCA performed in this analysis may be a strong predictor for expression among ncRNA. RNAP2 signals around ncRNA increased as PCA1 score decreased (Fig 4A). The same is true for miRNA and snRNA showing the same trend as expression (Fig 4C and 4D).

**Figure 3.**
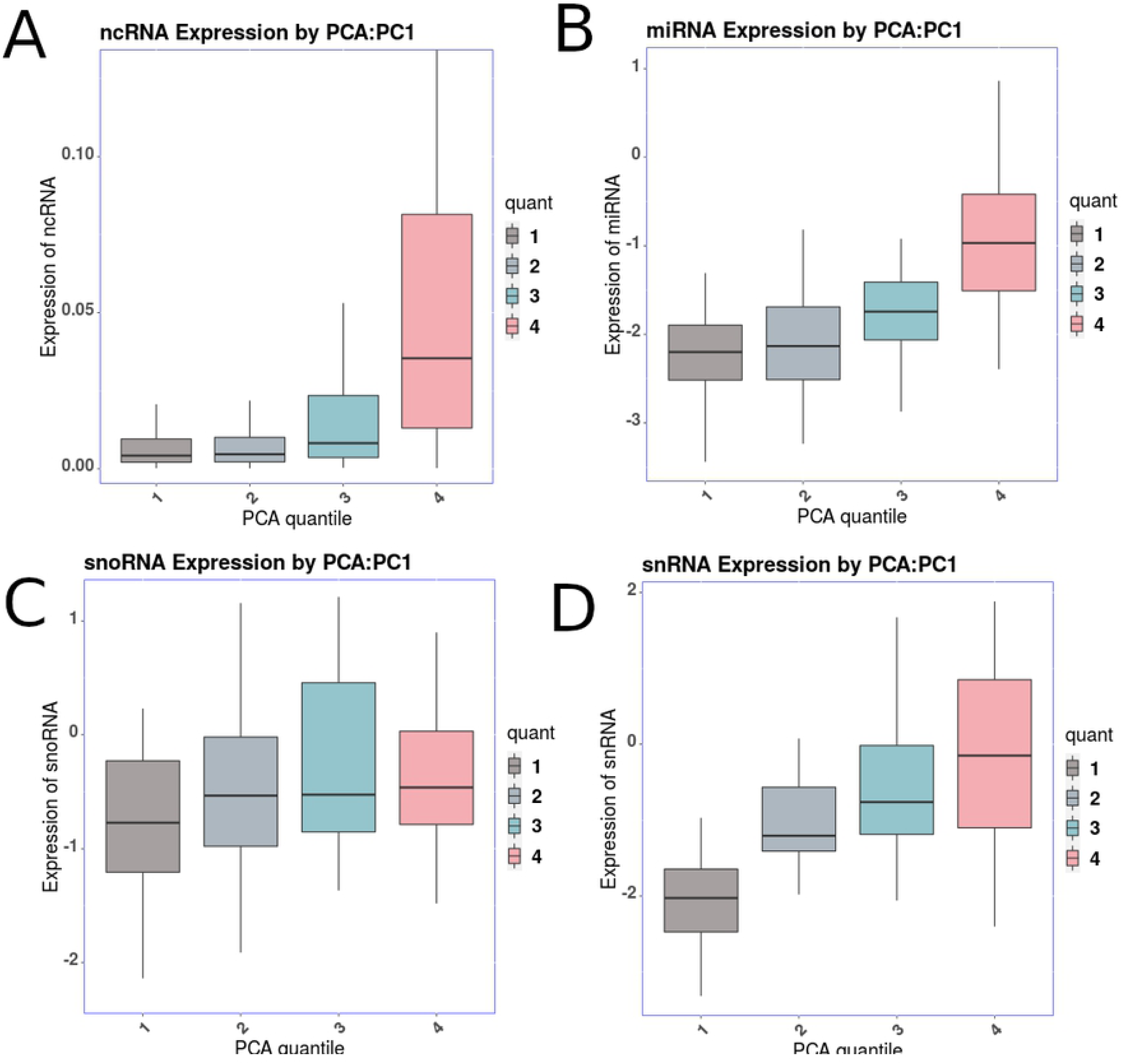
Box plot of expression for PCA1 quantile for ncRNA, miRNA, snoRNA and snRNA.

**Figure 4.**
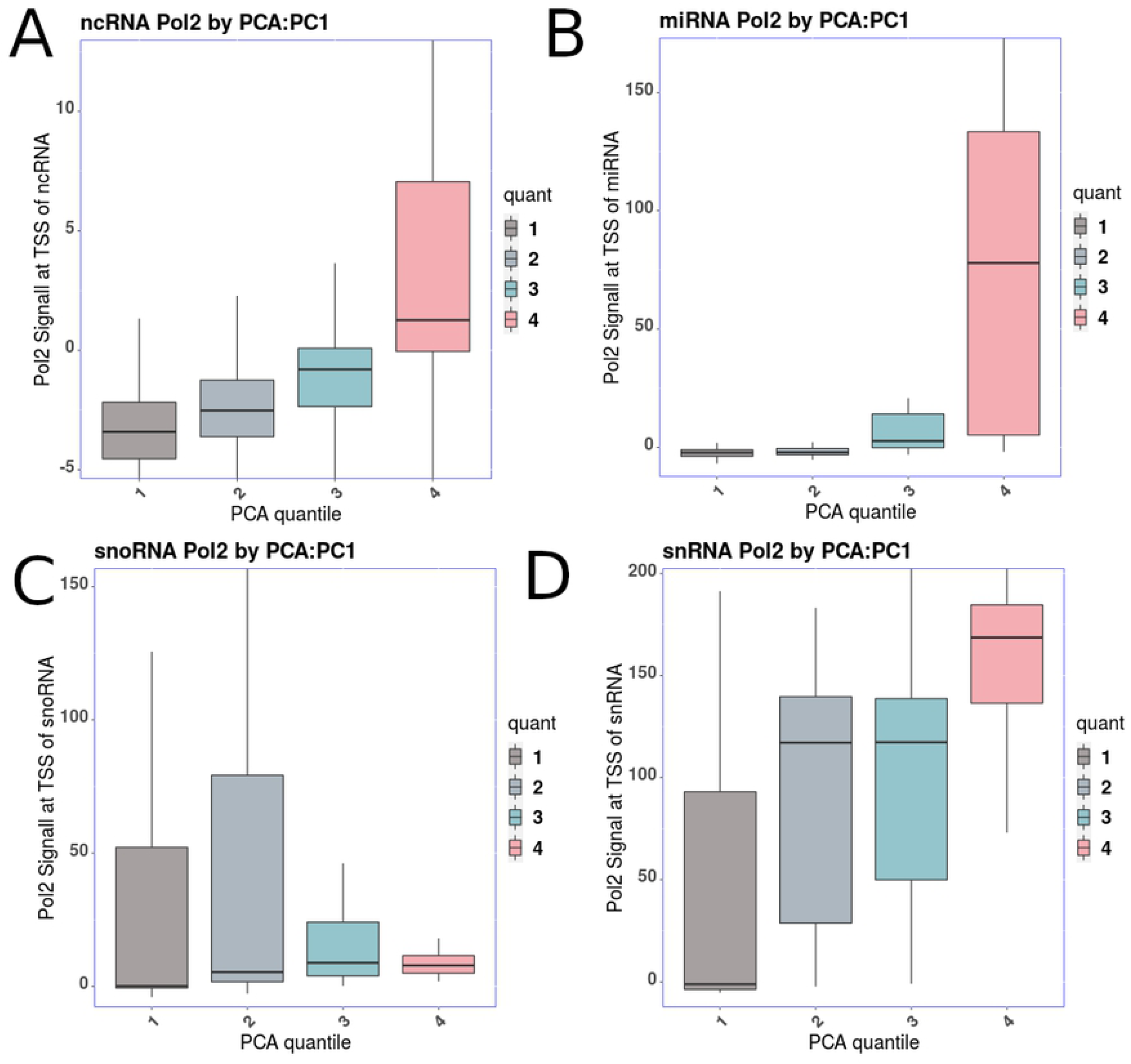
Box plot of RNA polymerase 2 binding at the TSS for PCA1 quantile for ncRNA, miRNA, snoRNA and snRNA.

Based on these observations, we believe that we have created a model of RNA polymerase recruitment based on histone modification. It has previously been shown that polymerase recruitment has a high correlation with expression of ncRNA (17). Therefore, the lower correlation we see between the PCA and expression is likely due to post transcriptional differences in ncRNA half-life due to degradation and sequestration. The half-life of ncRNA varies and may reflect the function of the ncRNA (18).

### Histone Modifications Signal Surrounding ncRNA Species

Having created a model of expression and RNAP2 binding, we next wanted to explore what histone modifications were enriched surrounding the TSS of the ncRNA species based on the PCA1 binning. Histone modification signals binned based on PCA1 had strong signals for ncRNA for the following histone modifications: H3K4me1, H3K4me2, H3K4me3, and H3K27ac (Fig 5A). All of these modifications have previously been shown to be enriched in some capacity in highly transcribed mRNAs. There is also a strong negative signal for H3K27me3 and the same group of ncRNA. This modification has been repeatedly shown to be correlated with repression of expression of mRNAs. This again shows a strong positive correlation between the PCA signal and the pol2 signal with the strength of the correlation decreasing among the other quantiles. This is a good confirmation of the correlation we saw previously (Fig 3).

**Figure 5.**
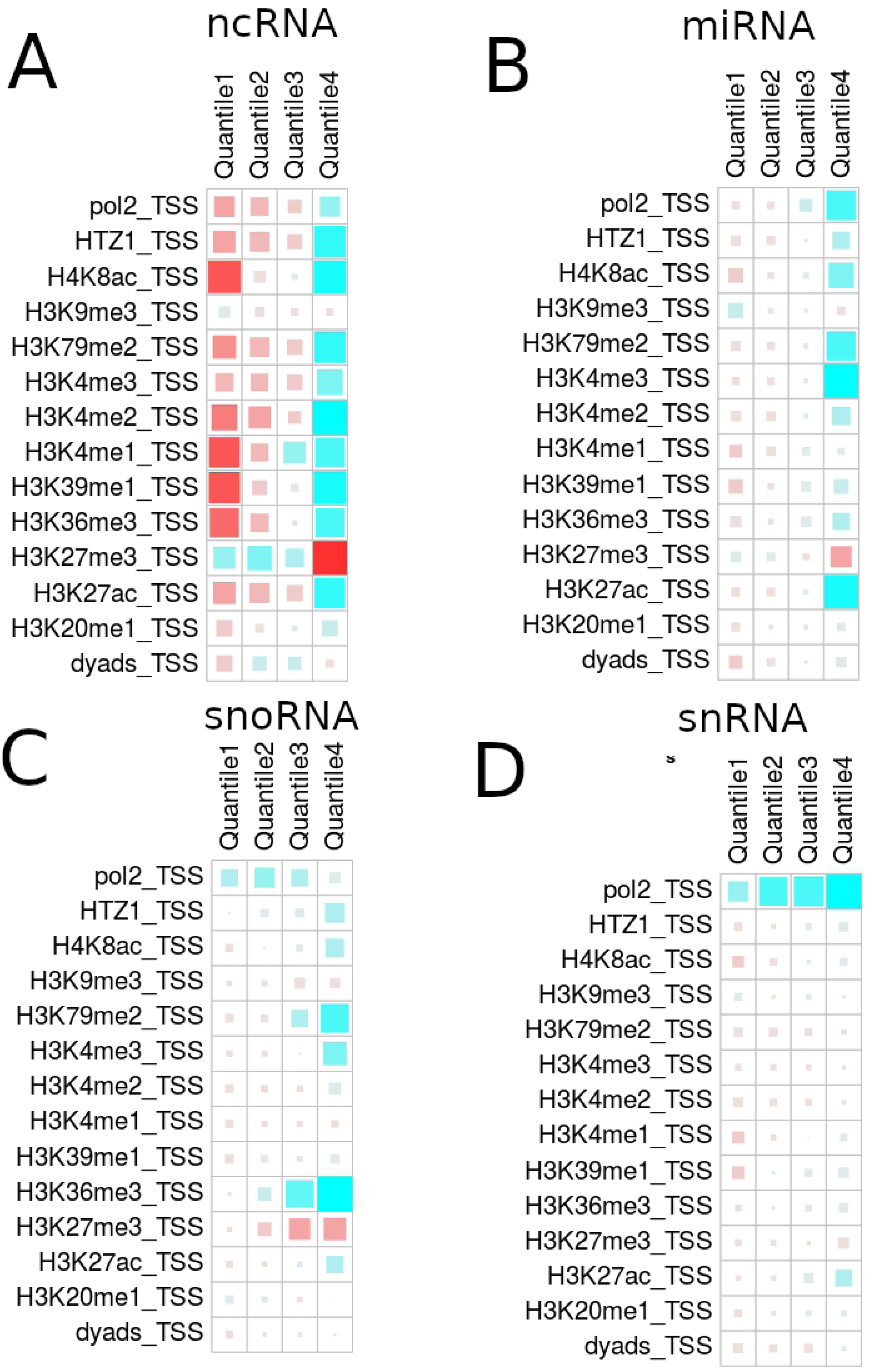
Histone modification ChIP-seq signal for each ncRNA (a), miRNA (b), snoRNA (c), and snRNA (d) by PCA quantile. Blue indicates a positive signal of that ChIP-seq in that quantile while red indicates a negative signal. The larger the square the larger the positive or negative signal for that ChIP-seq in that quantile.

A similar pattern is seen for miRNA. There are strong signals in the fourth quantile for H3K4me1, H3K4me2, H3K4me3, and H3K27ac and a strong negative signal in the fourth quartile and H3K27me3. A similar pattern exists among the snoRNAs (Fig 5C), however the signals are somewhat weaker for these modifications among the snoRNAs and the RNAP2 signal in the fourth quantile is weaker than the first three. The signal for RNAP2 among miRNA is weak in the first quantile and gradually increases among the other quantiles. A similar RNAp2 pattern exists among the snRNAs (Fig 5D), however the RNAP2 signal is fairly strong among all four quantiles.

We created a composite plot of those histones with the strongest signal at the TSS of miRNA, ncRNA, snoRNA and snRNA (Fig 6). RNAP2 is recruited immediately upstream of the TSS of ncRNAs and persists at a relatively strong signal across the gene body (Fig 6B). The pattern of histone modifications surrounding the TSS of the ncRNA resembles that of protein coding genes. miRNA, while still showing strong active histone modification enrichment, has a pattern of enrichment distinct from ncRNA. Histone regulation around snRNA resembled more closely the pattern seen for miRNA than ncRNA.

**Figure 6.**
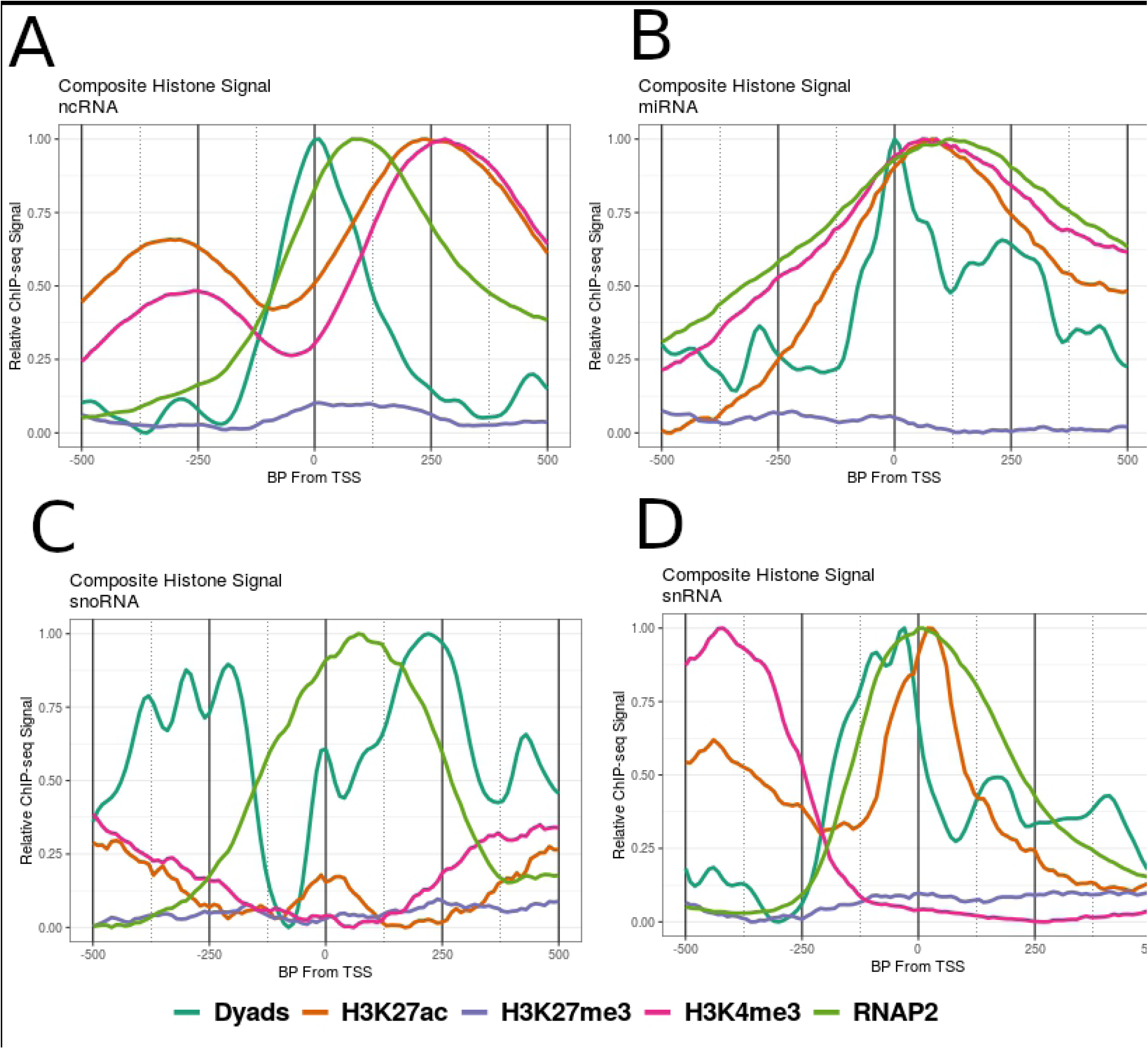
Chromosome Modification Composite plots for the PCA quantile with the highest expression profile for ncRNA, miRNA, snoRNA and snRNA.

## Discussion

Because mRNA has been the focus of the majority of studies focussed on histone modification regulation and because polyA enrichment of mRNA is an easy method for removing rRNA, regulation of expression of ncRNAs is less well understood. Further, analysis of regulation by histone modifications have similarly targeted at understanding mRNA regulation. The PCA and other analyses performed in this report show distinct patterns and roles for histone modifications in regulating different ncRNA species’ expression.

Our PCA shows an ability to predict expression for most ncRNA species except for tRNA. The lack of predictive ability for tRNAs was somewhat expected, given their repetitive nature in the genome and unique histone signatures in a number of organisms (19). Chromatin around tRNA genes is generally open which may help with steady-state transcription (20). Our correlation data show that there are strong correlations between our PCA results and between the RNAP2 signal which is correlated with expression levels of ncRNA species. This indicates that both our PCA and the RNAP2 signal around ncRNA can predict expression to a relatively high degree. We believe the deviations of ncRNA expression from this correlation are likely due to known post-transcriptional modifications of ncRNA species; therefore, we have created a model that predicts transcriptional activation, but its ability to fully predict how much RNA is present is masked by these post-transcriptional modifications and sequesterations.

We also show that the pattern of histone modifications surrounding actively transcribed ncRNA species differ by species. ncRNAs more closely resemble actively transcribed protein coding genes. However, miRNAs have a much different pattern despite enrichment of similar histone modifications. This suggests they may have a unique mechanism of activation. Interestingly, snRNA show a signal of MNase upstream of their annotated TSS, this may indicate that the actual snRNA TSS are actually upstream of their annotated TSS. This may be due to *C. elegans* unique splice leader events during transcription. Splice leader sequences are attached to the 5’ end of transcripts and allow an RNA transcript to create a number of unique mature transcripts (21). This type of analysis could indicate a novel method of predicting ncRNA output based on changes in histone modification signal.

As a result of this analysis, we confirmed that RNAP2 binding predicts RNA production, as would be expected based on the role RNAP2 plays in transcription. For some of the ncRNA species, the PCA model predicts RNAP2 binding better than any single histone modification. Based on this result and compared to results from other studies, we feel confident that RNAP2 is a good predictor of ncRNA expression. It remains an open question whether these observations are unique to this particular stage of C. elegans development, or if this is indicative of a broader trend.

